# Disrupting female flight in the vector *Aedes aegypti*

**DOI:** 10.1101/862300

**Authors:** Sarah O’Leary, Zach N. Adelman

## Abstract

*Aedes aegypti* is a vector of dengue, chikungunya, and Zika viruses. Current vector control strategies such as community engagement, source reduction, and insecticides have not been sufficient to prevent viral outbreaks. Thus, interest in novel strategies involving genetic engineering is growing. Female mosquitoes rely on flight to mate with males and obtain a bloodmeal from a host. We hypothesized that knockout of genes specifically expressed in female mosquitoes associated with the indirect flight muscles would result in a flightless female mosquito. With the CRISPR-Cas9 system, we performed embryonic microinjections of Cas9 protein and guide RNAs specific to genes hypothesized to control flight in mosquitoes, and have obtained genetic knockouts in several genes specifically expressed in the flight-muscle, including those specific to female flight muscle. Analysis of the phenotype of these female-specific gene knockout mutants resulted in flightless females and flying males. While further assessment is required, this work lays the groundwork for a mechanism of population control that is female-specific for the *Ae. aegypti* vector.

## Introduction

The yellow fever mosquito *Aedes aegypti* is a vector for many viruses of medical significance, such as dengue, Zika, chikungunya, and yellow fever. They can be found in tropical, subtropical, and temperate regions of the world, with outbreaks occurring in the US within the last century [1]. Only female *Ae. aegypti* bite to obtain a blood meal, which is required for egg production. After hatching from the embryo, *Ae. aegypti* like all other mosquito species will progress through the aquatic larval and pupal stages of their life, before emerging as an adult from the pupal casing to fly away [2, 3].

Currently, there is a safe and effective vaccine available for yellow fever [4], but not yet for dengue or Zika virus. Control efforts largely focus on reducing vector abundance, and include source reduction, and chemical methods like insecticides or larvicides [3, 5]. The short-term effect and high financial cost, along with the need for trained staff, presents challenges to the implementation, scaling, and maintenance of these control methods. With chemical methods, additional concerns relating to the emergence of resistance and effects on off-target species are increasing [6]. Because of these limitations, the need for novel vector control strategies is growing.

Genetic control strategies are receiving an increased amount of attention as viable vector control approaches, and include sterile insect technique (SIT) [7-9], release of an insect carrying a dominant lethal (RIDL) [7, 10], and gene drive. Population suppression approaches to vector control with genetic modifications seek to, in some way, prevent the female mosquito from being able to bloodfeed or mate, thus producing fewer or no offspring, leading to a population decline or collapse. Gene drive involves the spread of a genetic element beyond Mendelian rates of inheritance [11, 12]. Synthetic gene drive mechanisms can take advantage of the CRISPR/Cas9 system, which allows targeting of the genome at a precise location to catalyze a double-stranded break. Based on the type of DNA repair that occurs (non-homologous end joining or homology directed repair), one can achieve targeted disruption or insertion of a cargo sequence. Much work is being put in to understanding the formation of alleles resistant to CRISPR/Cas9 cleavage [13-15] and to increasing gene drive efficiencies overall [16-18]. Meanwhile, the CRISPR/Cas9 system has now become an efficient and inexpensive method for genome editing [19], and it has been utilized effectively in *Ae. aegypti* [20-23].

A function that is critical for both reproduction and survival in females is flight, as flight is required for mating, obtaining a blood meal, and escaping from water after eclosion. *AeAct-4* was identified previously as a female- and pupal-specific gene, with expression in the indirect flight muscles [24]. Other work has identified a male-specific actin gene [25] and male-specific myosin gene [26] related to flight, offering up the possibility that male and female flight in *Aedes* mosquitoes is controlled by these sex-specific genes.

To demonstrate the importance of female-specific actin and myosin genes to *Ae. aegypti* female flight, we used CRISPR/Cas9 to generate heritable loss-of-function alleles in *AeAct-4* and the female-specific myosin gene, along with a third gene, *Aeflightin*, expressed in both males and females. Phenotypic analysis of individuals homozygous for each introduced mutation in *AeAct-4* or the female-specific myosin confirmed that flight defects were both complete and likely restricted to females. Disruption of *Aeflightin* was associated with loss of flight in both sexes. The data support the pursuit of novel genetic strategies geared specifically for disrupting female flight in *Ae. aegypti*.

## Materials and methods

### Insect rearing

The Liverpool strain of *Ae. aegypti* was used for embryonic microinjections and outcrossing of mutant individuals. All mosquitoes were reared at 28°C and 80-85% humidity, with a 14/10 h light/dark light cycle. Ground up fish food (Tetra, Blacksburg, VA) was supplied throughout the aquatic developmental stages, and a cotton ball soaked with 10% sucrose solution was supplied during the adult stage. Flightless mosquitoes were supplied with raisins as the source of sucrose. Defibrinated sheep blood (Colorado Serum Company, Denver, CO) was offered for blood feeding, via a parafilm membrane feeder. Videos were taken with a Canon Rebel T3i digital camera.

### Guide RNA design and synthesis

Guide RNAs were designed by hand with the DNASTAR SeqBuilder Pro software (Madison, WI), using the appropriate gene sequence acquired through VectorBase. Primers used to make each sgRNA were ordered through the IDT custom DNA oligos web ordering service. Guide RNA synthesis was performed as previously described [20]. Briefly, Q5 High-Fidelity DNA Polymerase (New England BioLabs Inc., Ipswich, MA) was used for the PCR reaction, followed by the NucleoSpin Gel and PCR Clean-Up kit protocol (Machery-Nagel, Bethlehem, PA), the MEGAscript T7 Transcription kit protocol, and the MEGAclear Transcription Clean-Up kit protocol (Thermo Fisher Scientific, Waltham, MA). All sgRNAs were quantified, aliquoted, and stored at - 80°C. A list of oligos used to make each sgRNA are listed in table 2.

**Table 1.**
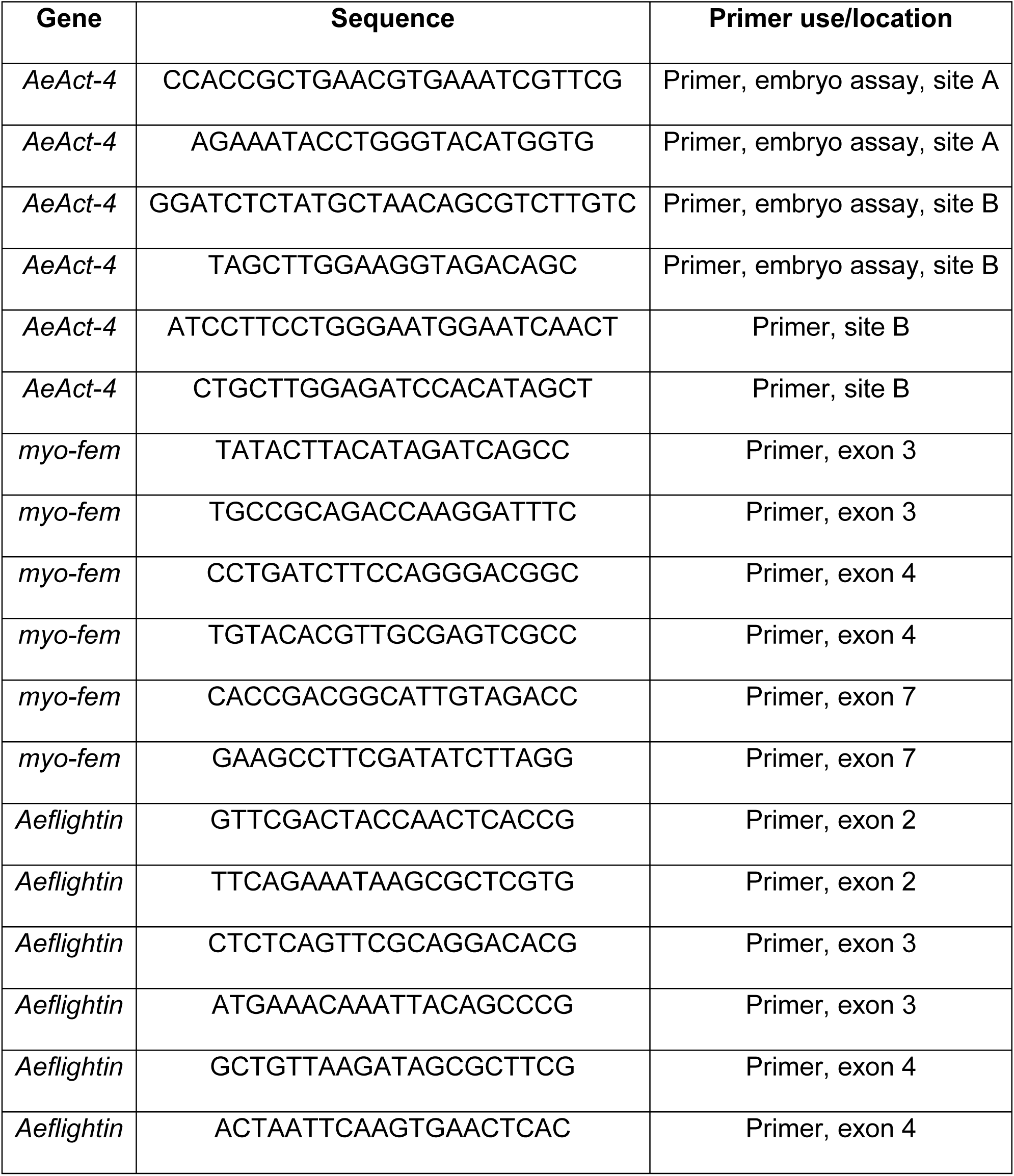
Primer oligo sequences.

**Table 2.**
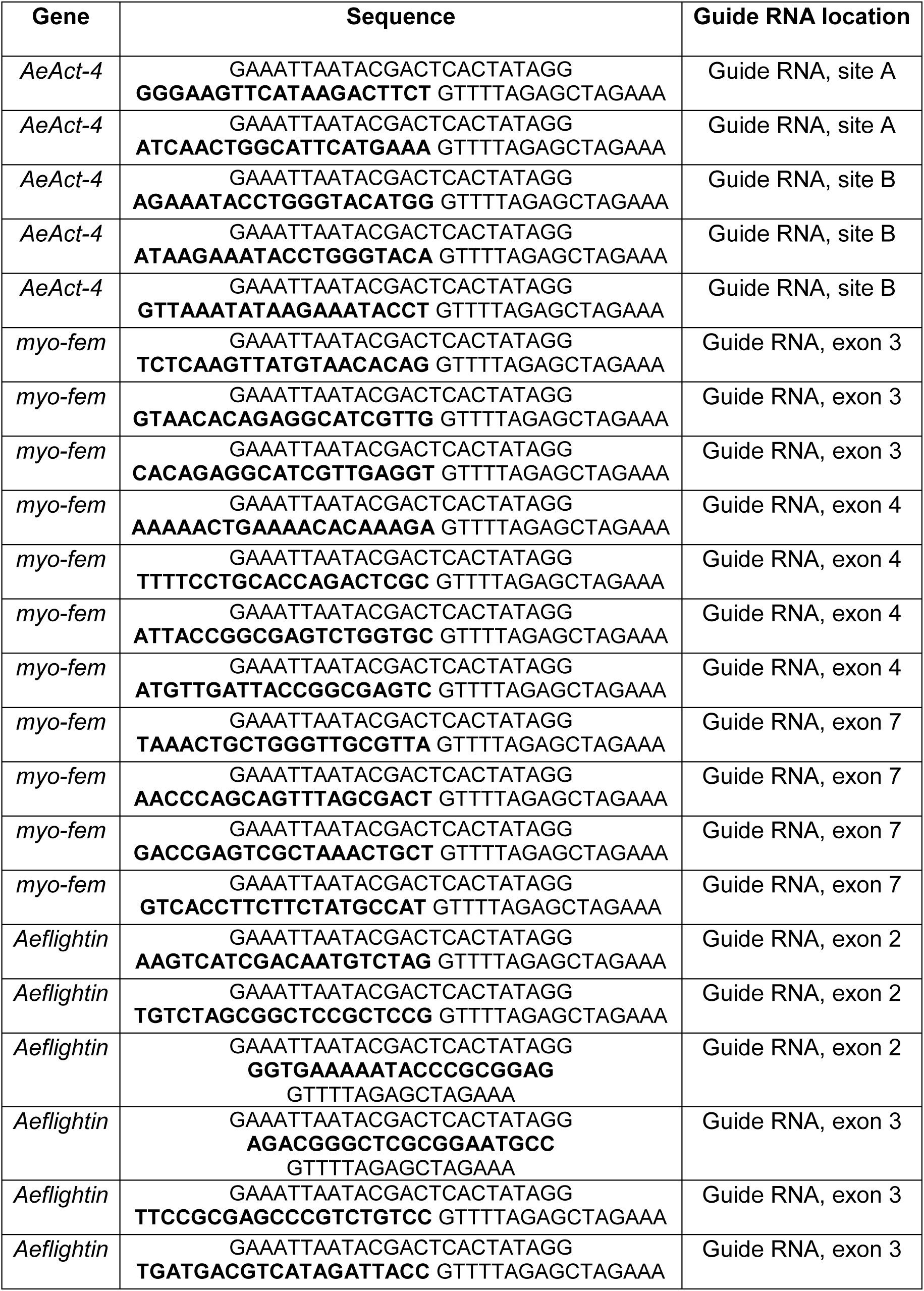
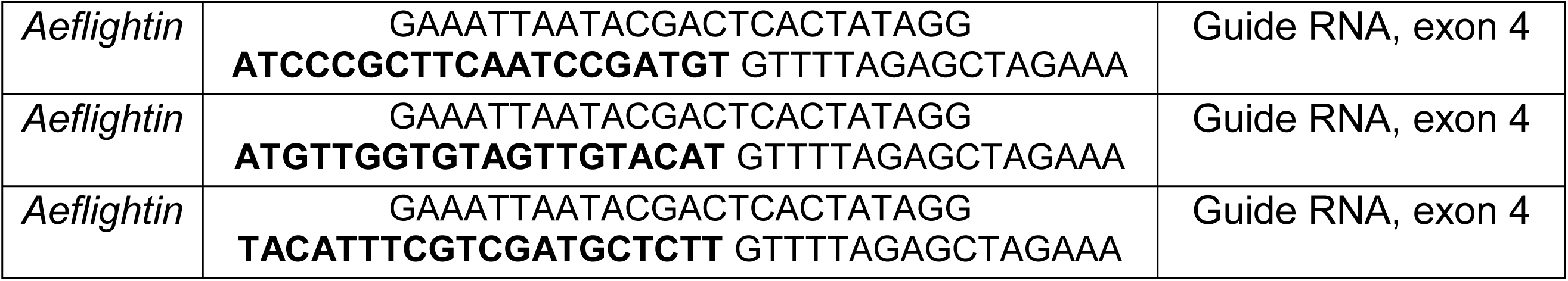
Guide RNA oligo sequences.

### Embryo microinjections

Borosilicate glass capillaries (World Precision Instruments Inc., Sarasota, FL) were pulled and beveled using the Sutter P-2000 Micropipette Puller and Sutter BV-10 Micropipette Beveller (Sutter Instrument Co., Novato, CA). Embryo microinjections were performed using a Leica DM 1000 LED Micromanipulator (Leica Biosystems, Buffalo Grove, IL) and FemptoJet 4i Microinjector (Eppendorf, Hauppauge, NY). Purified Cas9 protein (400 ng/μl) (PNA Bio, Thousand Oaks, CA) and sgRNAs (100 ng/μl) were combined into injection mixes, incubated at 37°C for 30 minutes, and centrifuged at max speed, 4°C, for a minimum of 45 minutes. Injections were performed into the posterior end of embryos that were less than three hours old. Injected embryos were either harvested at 24 hours (for embryo assays) or hatched after five days. Hatched G0 survivors were outcrossed to the wild type Liverpool strain.

### DNA extraction, PCR, HRMA, and Sanger sequencing

Genomic DNA was extracted from non-injected or sgRNA-injected Liverpool embryos following the Nucleospin Tissue kit protocol (Machery-Nagel, Bethlehem, PA). Embryo assays were performed with the LightScanner Master Mix kit (Idaho Technology Inc., Salt Lake City, UT), and mutant detection was performed on adult legs with the Phire Animal Tissue Direct PCR kit (Thermo Fisher Scientific, Waltham, MA) with added LCGreen Plus+ Melting dye (Idaho Technology Inc., Salt Lake City, UT). All samples were amplified with the C1000 Touch Thermal Cycler (Bio-rad, Hercules, CA) before being analyzed with the LightScanner Call-IT 2.0 software on the LightScanner instrument (Idaho Technology Inc., Salt Lake City, UT). Sanger sequencing was performed at the Laboratory for Genomic Technologies (Institute for Plant Genomics and Biotechnology, Texas A&M University, College Station, TX) and chromatograms were analyzed using Chromas software (Technelysium, Australia). A list of primers used are located in table 1.

### Flight determination

To assess flight, pupae were placed in plastic ketchup containers (with the stick of a Q-tip to aid in escaping from the water) in 5-quart plastic buckets (Home Depot) designed with a mesh covering over the top and a sock for internal access on the side. The plastic lining of the bucket inhibited the mosquitoes from climbing up the sides towards the mesh covering (adapted from Chae *et al*, “CRISPR/Cas9-driven Gene Knockout of *Aedes aegypti* Pupae-specific *Stretchin* Resulted in Flightless Mosquitoes”, in preparation). Flying mosquitoes that could fly up to the mesh covering were removed from the plastic bucket, and 24 hours was allowed to pass after the last pupae emerged before classifying the remaining mosquitoes as flightless. Dead adults whose flight phenotype could not be confirmed were not analyzed for a genotype.

## Results

Since female mosquitoes rely on flight to obtain a blood meal and mate, disrupting female-specific flight would prevent females from reproducing, or transmitting viruses. AeAct-4 has been characterized as a female- and pupal-specific flight protein that is expressed in the indirect flight muscles [24]. Analysis of RNA-seq data from the developmental transcriptome of *Ae. aegypti* [27] led us to select two other hypothesized flight genes, based on female-specific expression or pupal-specific expression.

The first gene, AAEL005656, is a paralog to the gene *myo-sex* (AAEL021838), a male-specific myosin gene located in the M locus on chromosome one [26] and required for male flight [28]. As AAEL005656 was found to be expressed specifically in female pupae, we reasoned that it might be similarly critical for female flight, and refer to this gene as *myo-fem*. The second gene, AAEL004249, is expressed in both males and females, but is pupal-specific. In *Drosophila melanogaster*, the ortholog to AAEL004249, *flightin*, has been shown to be expressed in the indirect flight muscles [29], with knockout resulting in a loss of flight ability [30]. We reasoned that despite lacking female specificity, *Aeflightin* may be a good target for disrupting female flight, so long as a male-specific rescue can be performed.

For each of these three target genes, we designed multiple sgRNAs. Due to high nucleotide sequence similarity between *AeAct-4* and other actin paralogs, we performed a multiple sequence alignment prior to sgRNA design. Eleven paralogs with ≥60% nucleotide sequence identity to *AeAct-4* were aligned to identify PAM sites that were unique to *AeAct-4*. For *myo-fem*, guide RNAs were designed to target the motor domain to ensure disruption of myosin function. For *Aeflightin*, guide RNAs were designed to each exon as we did not identify any paralogous genes in the *Ae. aegypti* genome.

Following in vitro synthesis, sgRNAs were complexed with Cas9 protein and injected in groups of 3-4 into *Ae. aegypti* embryos, which were harvested after 24 hours. To identify those batches capable of inducing strong gene disruption, genomic DNA from injected embryos was used as a template for PCR/HRMA of the target region (figure 1). For *AeAct-4*, we identified one location that was disrupted by a cluster of guide RNAs. For *myo-fem* and *Aeflightin*, we identified two clusters of guide RNAs with detectable editing in embryos.

**Figure 1.**
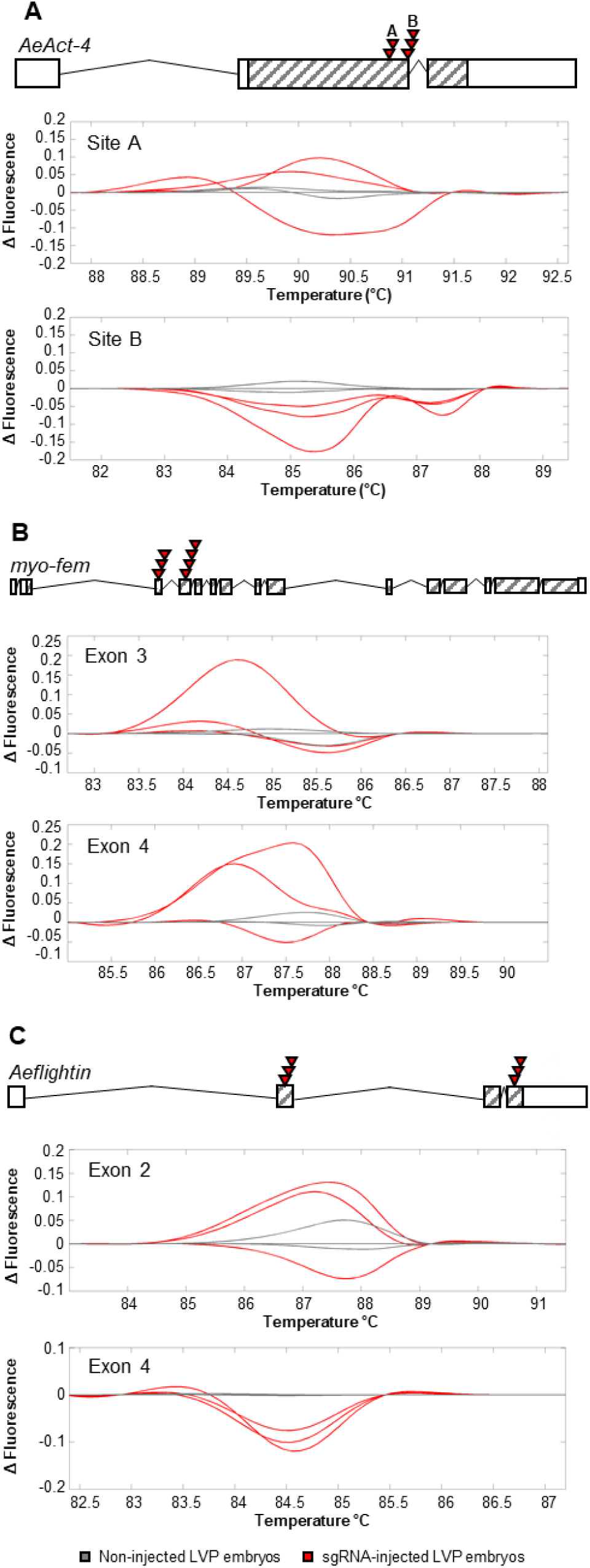
Validation of CRISPR reagents for *Ae. aegypti* genes involved in flight. Models for each target gene (**A**, *AeAct-4*; **B**, *myo-fem*; **C**, *Aeflightin*). Boxes represent exons, cross-hatched areas represent the ORF of the corresponding mRNA. For each, red triangles indicate the locations of sgRNAs that were found to successfully cleave the DNA during embryo assay. Melt curves are displayed for each location of sgRNAs (indicated as “site” or “exon”), with sgRNA-injected (red) and non-injected (gray) samples.

For each mixture of guide RNAs that displayed editing activity, we repeated embryo microinjections and this time allowed the embryos to hatch after five days. Survivors were outcrossed to the parental Liverpool strain to obtain G1 progeny. G1 adults were screened via PCR/HRMA and Sanger sequencing for out-of-frame mutations (table 3). Deletions predicted to result in a frameshift mutation were recovered for *AeAct-4, myo-fem*, and *Aeflightin* (figure 2). Genotyped adults with out-of-frame mutations were outcrossed to the parental strain for two generations followed by an intercross of heterozygous individuals (figure 3). Since we were targeting female-specific genes, we sought to observe the ratio of males to females during development to ensure there was no early mortality associated with either sex. For knockout of all three of our target genes, the observed male to female ratios were not significantly different from the expected 1:1 ratio (*AeAct-4* p = 0.8124; *myo-fem* p = 0.9390; *Aeflightin* p = 0.0828, Chi square analysis). This indicates there was no significant female- or male-associated mortality prior to the adult stage associated with knockout of these genes.

**Table 3.** Generation of loss-of-functions mutants in *Ae. aegypti* flight genes using CRISPR/Cas9.

| Line | #<br>Injected | # G<sub>0</sub><br>Hatched | % G<sub>0</sub><br>Survival | # G<sub>1</sub><br>Genotyped | # G<sub>1</sub><br>Sequenced | # G<sub>1</sub><br>Mutants | Results |
| --- | --- | --- | --- | --- | --- | --- | --- |
| *AeAct-4* Exp. 1* | 407 | 83 | 20.4% | 240 | 19 | 2 | Δ3 |
| *myo-fem* Exon 3<br>Exp. 1 | 468 | 43 | 9.2% | 320 | 20 | 9 | Δ4Δ4, Δ3, i2, Δ11 |
| *myo-fem* Exon 4<br>Exp. 1 | 455 | 54 | 11.9% | 197 | 6 | 0 | N/A |
| *Aeflightin* Exon 2<br>Exp. 1 | 373 | 49 | 13.1% | 80 | 10 | 2 | Δ9 |
| *Aeflightin* Exon 4<br>Exp. 1 | 353 | 57 | 16.2% | 240 | 4 | 2 | Δ3 |
| *AeAct-4* Exp. 2* | 344 | 120 | 34.9% | 0 | 0 | 0 | N/A |
| *myo-fem* Exon 4<br>Exp. 2 | 265 | 29 | 11.0% | 80 | 0 | 0 | N/A |
| *Aeflightin* Exon 2<br>Exp. 2 | 362 | 114 | 31.5% | 240 | 23 | 15 | Δ2Δ5a, Δ5b, Δ4,<br>Δ2Δ5b, Δ5c |
| *Aeflightin* Exon 4<br>Exp. 2 | 268 | 58 | 21.6% | 80 | 0 | 0 | N/A |
| *AeAct-4* B Exp. 2* | 268 | 66 | 24.6% | 80 | 10 | 4 | Δ2, Δ10 |

\*“*AeAct-4* Exp. 1” injection mixes included *kmo* sgRNAs. “*AeAct-4* B Exp. 2” injection mixes included only site B sgRNAs, as opposed to site A and B for *AeAct-4*. “*AeAct-4* Exp. 2” G<sub>1</sub> individuals were not hatched and screened due to increased sequence variability in the amplicon used for HRMA.

**Figure 2.**
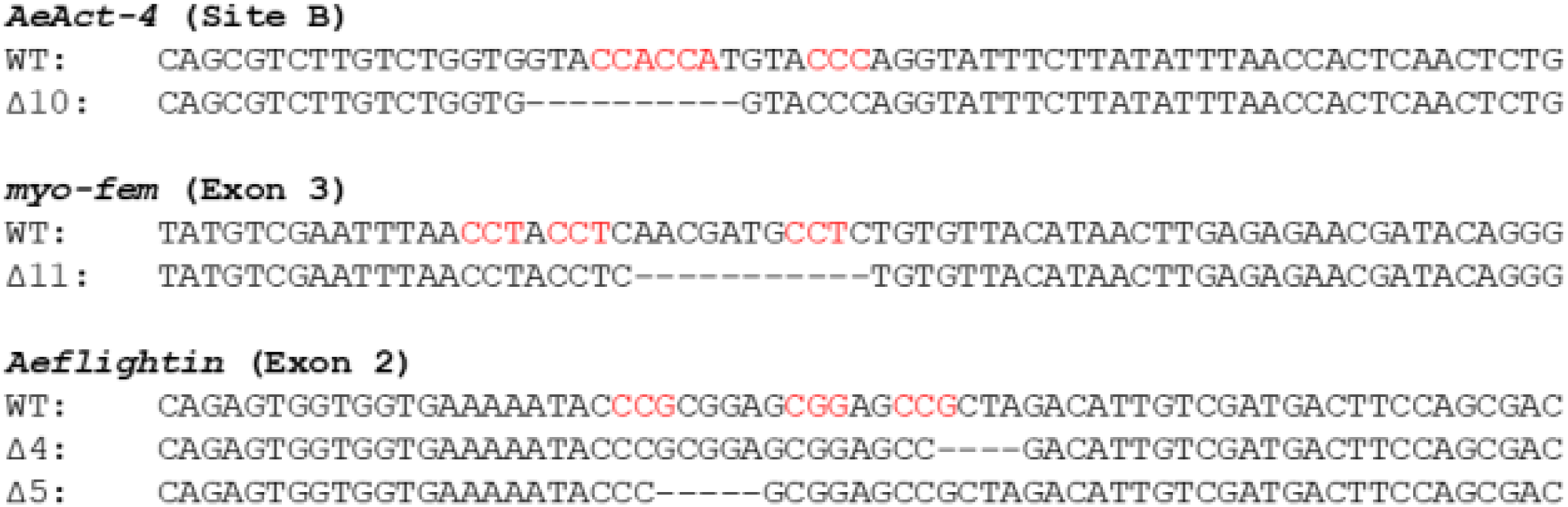
Heritable loss-of-function mutations in *Ae. aegypti* flight genes. For each gene, the wild type (WT) and mutant sequence is shown. The PAM sites for each sgRNA used in the injection mix for the specified location is highlighted in red.

**Figure 3.**
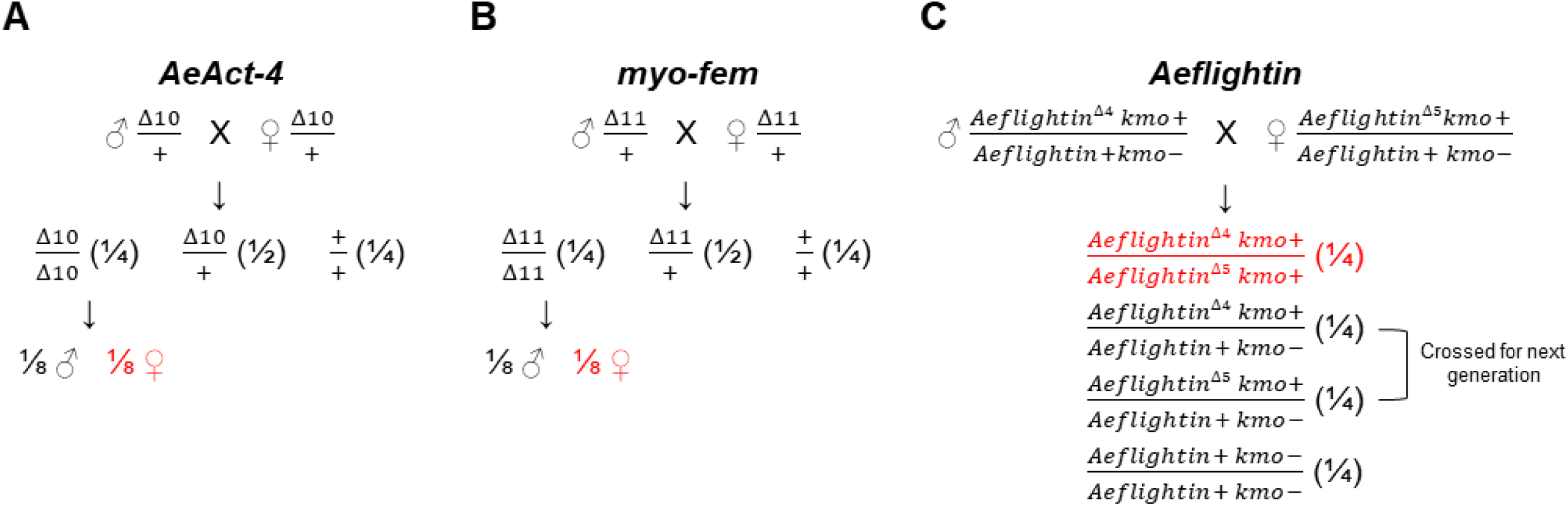
Expected genotypic and phenotypic ratios. A heterozygous self-cross for each gene (*AeAct-4* in A, *myo-fem* in B, and *Aeflightin* in C) is represented above. The ratio for each genotype is noted in parenthesis, with the individuals with an expected flightless phenotype highlighted in red. For *Aeflightin*, which was linked to *kmo* in a previous outcrossing, phenotypic identification of all white-eyed pupae enables removal before further phenotypic analysis based on flight (see figure 4) is performed.

To assess any potential phenotype related to flight of homozygous mutants (figure 4), G4 progeny were obtained following the workflow in figure 4 to characterize mosquitoes as flightless or flying. Upon stimulation of flight, control mosquitoes could fly, while some from each test cross could not; these were therefore categorized as flightless and hypothesized to be homozygous for each targeted gene disruption. Flightless individuals appeared to have various alternative wing phenotypes when resting that differed from wild type (figure 5). Some individuals could move or beat their wings, but with no success in flight. Other individuals could initiate flight takeoff, but not sustain flight. Agitation to provoke flight via shaking and tapping of the plastic buckets (video 1) or gentle spraying of condensed air were used to evaluate flight ability; flightless mosquitoes remained so regardless of the method of stimulation.

**Figure 4.**
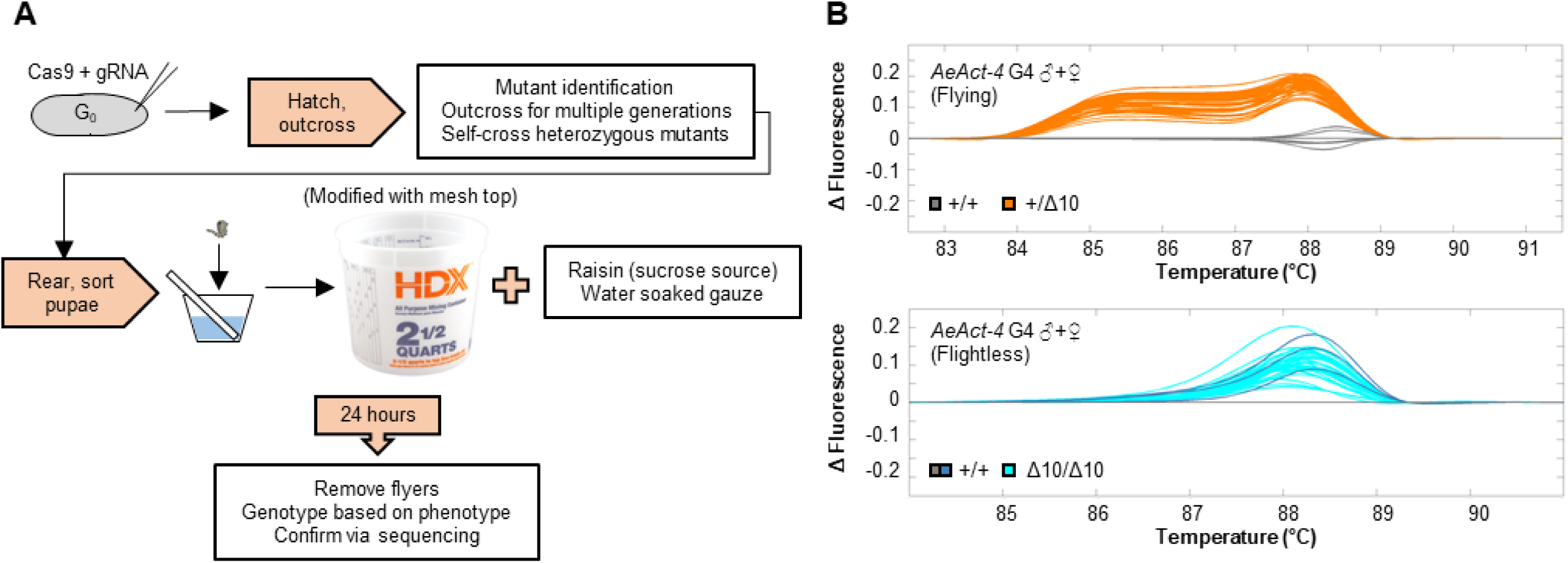
Separation of flightless from flying *Ae. aegypti*. (**A**) Workflow from initial injection of CRISPR/Cas9 reagents to analysis of flightless phenotypes and genotyping assays. (**B**) HRMA results for flying and flightless phenotypes associated with *AeAct-4*.

**Figure 5.**
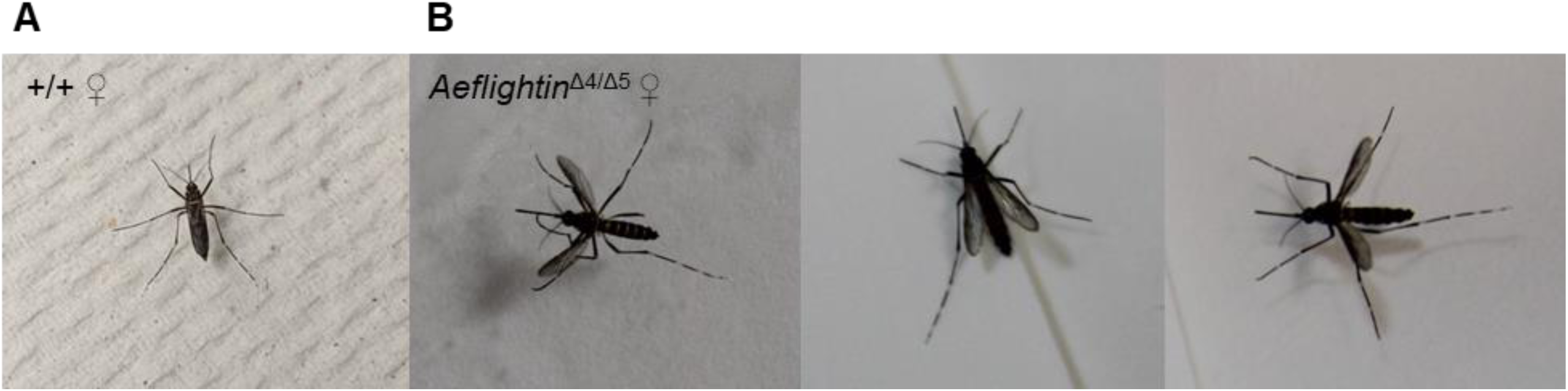
Disrupting of *Ae. aegypti* flight genes results in altered wing phenotypes. Wild type female (**A**) or *Aeflightin*^Δ4/Δ5^ females (**B**) when resting.

As all observations of flight behavior were made without consideration for genotype, we sought to determine whether there was a relationship between the ability of our mosquitoes to fly and inheriting one or two copies of each loss-of-function mutation. A genotypic analysis of all flightless individuals and a subset of flying individuals was performed for each cross (tables 4-6, figure 6). For both *AeAct-4* (table 4) and *myo-fem* (table 5), males homozygous for the loss-of-function mutations could fly (though for *AeAct-4* there was also a single homozygous male that could not fly). However, all homozygous *AeAct-4*^*Δ10*^ females were flightless (100%). Interestingly, while most heterozygous *AeAct-4*^*Δ10*^ females could fly (92%, only two were flightless), this was not the case for *myo-fem*^*Δ11*^, where most heterozygous (88%) and all homozygous (100%) females were flightless. The flying to flightless ratio did not differ significantly from the expected ratio for *AeAct-4* (p = 0.5300, Chi square analysis), suggesting the flightless phenotype was recessive in this case. Due to the increase in flightless heterozygous females for *myo-fem*, the flying to flightless ratio was significantly different from the expected ratio (p < 0.0001, Chi square analysis), confirming that defects in *myo-fem* are not recessive. As the *Aeflightin* gene is tightly linked to the *kmo* gene involved in eye pigmentation, *Aeflightin* mutants were outcrossed to a kynurenine 3-monooxygenase (*kmo*) knockout strain [20] to help track the corresponding genotypes. This aided in phenotypic identification of homozygous *kmo* individuals who do not carry the *Aeflightin* mutation, as well as maintenance of a transheterozygous line. At G4, low levels of recombination between *kmo* and *Aeflightin* were both expected and observed (table 6). The white- and black-eyed ratio did not differ significantly from the expected ratio (p = 0.4474, Chi square analysis) as the *kmo* loss-of-function is completely recessive. White-eyed adults hypothesized as wild type for *Aeflightin* based on HRMA results were confirmed via sequencing, while black-eyed adults that could fly were confirmed as heterozygous for either the *Aeflightin*^*Δ4*^ or *Aeflightin*^*Δ5*^ mutations, with only a small portion of heterozygous individuals (8%) that were flightless. The flying to flightless ratio did not differ significantly from the expected ratio for *Aeflightin* (p = 0.0008, Chi square analysis). Critically, all individuals confirmed as transheterozygous (*Aeflightin*^*Δ4/Δ5*^) were flightless, confirming that *Aeflightin* is required for flight in both males and female *Ae. aegypti*.

**Table 4.** Phenotypic and genotypic analysis of *AeAct-4* G4 individuals.

| ***AeAct-4* G3: ♂ $\Delta 10/+$ X ♀ $\Delta 10/+$** |  |  |  |  |  |
| --- | --- | --- | --- | --- | --- |
| **Phenotypic analysis** |  |  |  |  |  |
| **Male:**<br>140/284 (49.3%) |  |  | **Female:**<br>144/284 (50.7%) |  |  |
| **Flying:**<br>245/284 (86.3%) |  |  | **Flightless:**<br>39/284 (13.7%) |  |  |
| **Genotypic analysis** |  |  | **Genotypic analysis** |  |  |
| **$\Delta 10/\Delta 10$:** | **Male:**<br>4/39<br>(10.3%) | **Female:**<br>0 | **$\Delta 10/\Delta 10$:** | **Male:**<br>1/1<br>(100%) | **Female:**<br>36/38<br>(94.7%) |
| **$\Delta 10/+$:** | 22/39<br>(56.4%) | 24/40<br>(60.0%) | **$\Delta 10/+$:** | 0 | 2/38<br>(5.3%) |
| **$+/+$:** | 3/39 (7.7%) | 0/40 (0%) | **$+/+$:** | 0 | 0 |
| **Undetermined:** | 10/39<br>(25.6%)* | 16/40<br>(40.0%)* | **Undetermined:** | 0 | 0 |

\*Only a subset of individuals analyzed by HRMA was confirmed by sequencing.

**Table 5.** Phenotypic and genotypic analysis of *myo-fem* G4 individuals.

| ***myo-fem* G3: ♂ $\Delta 11/+$ X ♀ $\Delta 11/+$** |  |  |  |  |  |
| --- | --- | --- | --- | --- | --- |
| **Phenotypic analysis** |  |  |  |  |  |
| **Male:**<br>85/171 (49.7%) |  |  | **Female:**<br>86/171 (49.7%) |  |  |
| **Flying:**<br>112/171 (65.5%) |  |  | **Flightless:**<br>59/171 (34.5%) |  |  |
| **Genotypic analysis** |  |  | **Genotypic analysis** |  |  |
| **$\Delta 11/\Delta 11$:** | **Male:**<br>2/28 (7.1%) | **Female:**<br>0 | **$\Delta 11/\Delta 11$:** | **Male:**<br>0 | **Female:**<br>18/55<br>(32.7%) |
| **$\Delta 11/+$:** | 11/28<br>(39.3%) | 5/24<br>(20.8%) | **$\Delta 11/+$:** | 0 | 37/55<br>(67.3%) |
| **$+/+$:** | 4/28<br>(14.3%) | 0/24 (0%) | **$+/+$:** | 0 | 0 |
| **Undetermined:** | 11/28<br>(39.3%)* | 19/24<br>(79.2%)* | **Undetermined:** | 0 | 0 |

\*Only a subset of individuals analyzed by HRMA was confirmed by sequencing.

**Table 6.**
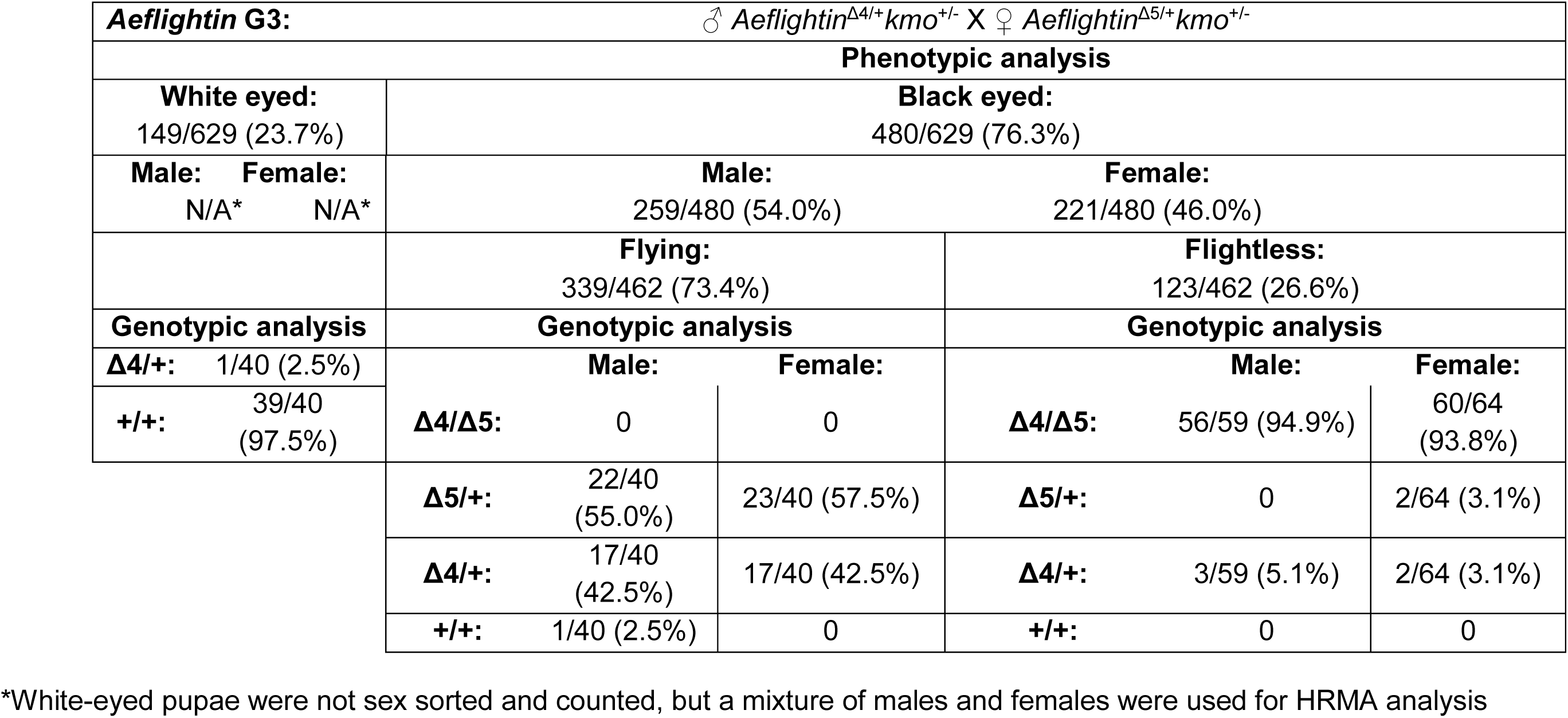
Phenotypic and genotypic analysis of *Aeflightin* G4 individuals.

**Figure 6.**
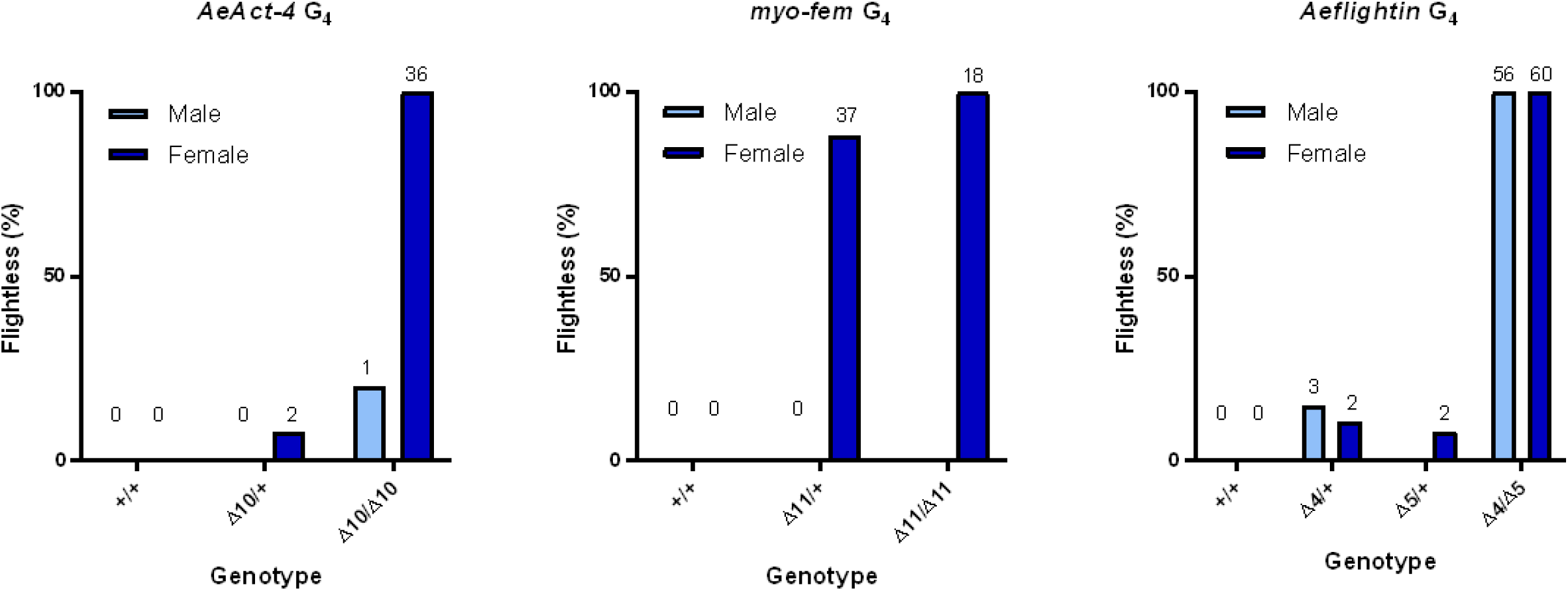
Flightless percentages based on genotypes for each gene. The percentage of flightless male or female mosquitoes for (**A**, *AeAct-4*; **B**, *myo-fem*; **C**, *Aeflightin*). The number above each bar represents the number of individuals displaying the flightless phenotype, and were confirmed for the specified genotype via sequencing.

## Discussion

Our results indicate successful knockout of three flight-specific genes, two of which are expressed predominantly in females (*AeAct-4* and *myo-fem*). Interestingly, we found that while *AeAct-4* and *Aeflightin* were haplosufficient, two intact copies of *myo-fem* appeared to be required for normal female flight. We note though that our approach allowed 24 hours after the last adult emerged for all mosquitoes to gain the ability to fly. For *myo-fem*, there were a few individuals who subsequently gained the ability to fly up to 48 hours after all flyers had been removed. These individuals seemed to have a delay (≥ 24 hours post-eclosion) in gaining flight ability, suggesting that a single copy of *myo-fem*, while insufficient to program the normal timing of development of the flight muscles, may be sufficient provided the female can survive long enough. If *myo-fem* is truly haploinsufficient, this opens the door for development of strong synthetic sex distorters for suppressing *Ae. aegypti* populations. Despite the lack of female specificity for the third gene, *Aeflightin*, we reason that this still represents a useful target so long as a male-specific rescue can be performed to fully restore male flight.

Flightless *Ae. aegypti* have been developed previously through the transgenic overexpression of the tTa transactivator specifically in the female flight muscle [31]. In this case, the promoter region of *AeAct-4* was used to control transgene expression, however transgenic males were found to have decreased mating competitiveness in field cage trials [32, 33]. Variability in the level of transgene overexpression also resulted in incomplete penetrance of the flightless phenotype. In our case, disruption of the *AeAct-4* gene through heritable gene editing resulted in a completely penetrant phenotype without the requirement for continuous transgene expression. However, it will be important to assess the fitness of the males homozygous for either the *AeAct-4* or *myo-fem* mutations, as both *AeAct-4* and *myo-fem* show low levels of transcription in male pupae [27], and so it is possible that male flight, while not disrupted, could be reduced in these individuals when more intensive flying is required.

Disrupting flight specifically in female mosquitoes could be used to achieve sex distortion of the adult population. This is conceptually similar to other sex distortion approaches such as the X-shredding system based on the I-PpoI homing endonuclease when active only during male meiosis, and shown to be capable of producing >95% male progeny [34]. Other examples of targets for sex-ratio distortion in *An. gambiae* include female reproductive genes [35, 36] and the female transcript of *doublesex*, which causes an intersex phenotype and complete sterility [37], however the recessive nature of these phenotypes reduces their power as sex distorters. In *Drosophila melanogaster*, disruption of other genes causing female fertility or embryonic lethality [17] have been shown to skew the sex-ratio towards males, while in *Ae. aegypti* overexpression of the male sex determining factor *nix* has been proposed as a method of sex distortion [38]. We note that the development of a sex-ratio distortion approach that targets female-specific flight would allow for maximum competition for resources during larval development as well as allow active monitoring of the number of females doomed to flightlessness, while at the same time preventing the adult female from reproducing and potentially transmitting deadly pathogens. With the data presented here and with further analysis, we can begin to consider or develop an approach for a mechanism of population control that targets female-specific flight for the *Ae. aegypti* vector.

## Acknowledgements

We thank members of the Adelman lab for assistance in rearing mosquitoes, in particular Dr. Keun Chae for assistance in developing flight assay protocols. This work was supported by Texas A&M Agrilife Research (Insect Vectored Disease Grant Program) and the Genetics Program at Texas A&M University.

## Supplementary Figure Legends

**<u>Video 1. Flight tests.</u>** Flightless *AeAct-4* individuals on the left, and control individuals on the right. The plastic buckets used to contain the adult mosquitoes was agitated by knocking on either side.

